# CXCR4^+^ Treg cells control serum IgM levels and natural IgM autoantibody production by B1 cells in the bone marrow

**DOI:** 10.1101/2021.11.14.468515

**Authors:** Shlomo Elias, Rahul Sharma, Michael Schizas, Izabella Valdez, Sun-Mi Park, Paula Gonzalez-Figueroa, Quan-Zhen Li, Beatrice Hoyos, Alexander Y. Rudensky

**Affiliations:** Immunology Program and Memorial Sloan Kettering Cancer Center, New York, NY, USA; Howard Hughes Medical Institute, and Memorial Sloan Kettering Cancer Center, New York, NY, USA; Molecular Pharmacology Program, Memorial Sloan Kettering Cancer Center, New York, NY, USA; Dept of Immunology and Infectious Disease, The John Curtin School of Medical Research, The Australian National University, Canberra, Australia; Microarray and Immune Phenotyping Core Facility, Department of Immunology, University of Texas Southwestern Medical Center, Dallas, TX, USA

**Keywords:** Treg cells, CXCR4, bone marrow, B-1 B cells, IgM

## Abstract

Regulatory T (Treg) cells represent a specialized lineage of suppressive CD4^+^ T cells, whose functionality is critically dependent on their ability to migrate to, and dwell in the proximity of cells they control. Here we show that continuous expression of the chemokine receptor CXCR4 in Treg cells is required for their ability to accumulate in the bone marrow (BM). Induced CXCR4 ablation in Treg cells led to their rapid depletion and consequent increase in mature B cells, foremost B-1 subset, observed exclusively in the BM without detectable changes in plasma cells or hematopoietic stem cells or any signs of systemic or local immune activation elsewhere. Dysregulation of BM B-1 B cells was associated with a highly specific increase in IgM autoantibodies and the total serum IgM levels. Thus, Treg cells control autoreactive B-1 B cells in a CXCR4-dependent manner. These findings have significant implications for understanding of regulation of B cell autoreactivity and malignancies.

## Introduction

Cells of the immune system exhibit mixed mobile and sedentary lifestyles to confer protection of an organism against a wide range of extrinsic biotic and abiotic challenges and intrinsic perturbations of organismal homeostasis. Chemokine receptors play critical roles in enabling migration of precursors and recirculation of mature immune cells through lymphoid and non-lymphoid organs as well as in their dynamic positioning within these tissues. Shared expression of chemokine receptors with the same ligand specificity by different immune cell types facilitates their encounters. These temporally and spatially coordinated interactions are paramount for the elaboration of immune responses and their regulation. Regulatory CD4^+^ T (Treg) cells, expressing transcription factor Foxp3, represent a specialized lineage which restrains responses of other immune cells (Fontenot et al., 2003; Hori et al., 2003; Sakaguchi et al., 1995). Immunosuppressive and tissue supporting functions of activated Treg cells are thought to require a close opposition to their target cells with matching chemokine receptor expression. Accordingly, Treg cells express receptors for a number of pro-inflammatory chemokines in addition to homeostatic secondary lymphoid organ homing receptor CCR7. Indeed, restricted CCR4 deficiency in Treg cells impaired their ability to control skin and lung inflammation (Annunziato et al., 2002; Huehn et al., 2004; Iellem et al., 2003; Sather et al., 2007).

In addition to CCR7 and pro-inflammatory chemokine receptors, Treg cells also express CXCR4, a chemokine receptor playing an important role in thymocyte differentiation and maturation, neutrophil and B cell retention in, and their release from the bone marrow (BM) and in neutrophil BM migration (Nie et al., 2004; Suratt et al., 2004). It is noteworthy that in the thymus, in addition to guiding migration of immature thymocytes, CXCR4 signaling, in cooperation with other receptors, can promote survival of T cells. Consistent with CXCR4 function in other immune cell types, Treg cells were shown to migrate to and accumulate in the BM, where they comprise a higher proportion – up to 40-50% - of the overall CD4 T cell population than in the majority of other lymphoid and non-lymphoid tissues (Hirata et al., 2018; Zou et al., 2004). BM Treg cells exhibited increased suppressive capacity *in vitro* compared to their peripheral blood counterparts, and displayed distinct gene expression features in comparison to splenic Treg cells (Camacho et al., 2020; Glatman Zaretsky et al., 2017; Zou *et al.*, 2004). Functional studies suggested that BM Treg cells support immune privileged status of the hematopoietic stem cell (HSC)-niche consistent with their proximity to the endosteal surface adjacent to HSCs and that Treg cell derived IL-10 facilitates HSC supporting function of BM stromal cells (Camacho *et al.*, 2020). BM-focused depletion of Treg cells upon selective ablation of CXCR4 or wholesale loss of Treg cells by administration of diphtheria toxin (DT) to *Foxp3^DTR^* mice were suggested to increase HSC numbers and their *in vitro* colony forming capacity (Hirata *et al.*, 2018; Pierini et al., 2017). Systemic ablation of Treg cells was also reported to decrease numbers of B-lineage cells in the BM across different maturation stages including pro-, pre- and mature B cells (Pierini *et al.*, 2017). Besides proposed support for HSC maintenance and B cell differentiation, BM Treg cells, which co-localize with CD11c^+^ cells and plasma cells in the BM, are thought to support the maintenance of the latter (Glatman Zaretsky *et al.*, 2017).

These observations suggest that expression of CXCR4 by Treg cells allows them to exert broadly targeted tissue-supporting accessory rather than immunosuppressive function in the BM, with the exception of allogeneic BM transplantation settings. This notion is confounded, however, by particularities of experimental setups of prior studies. The latter include generalized inflammatory responses and profound changes of organismal physiology caused by ablation of Treg cells in *Foxp3^DTR^* mice, uncertain degree of CD25 antibody mediated depletion of immature and mature cell types other than Treg cells and potential changes in Treg cell physiology due to CXCR4 ablation in Treg cells during their thymic differentiation by a constitutively expressed Cre recombinase encoded by the *Foxp3* locus (Lee et al., 2007). Thus, the specific roles of CXCR4 expression by Treg cells and function of CXCR4 expressing Treg cells remain poorly understood.

To gain insights into putative roles of CXCR4-expressing Treg cells in the BM we combined their extensive characterization with tamoxifen-inducible ablation of CXCR4 in Treg cells to specifically account for their CXCR4-dependent functions. Upon temporally controlled selective depletion of Treg cells in the adult BM induced by tamoxifen-triggered CXCR4 loss, we observed an increase in BM resident, but not peritoneal B1-B cells in the BM, but not in the periphery. These increases were associated with a marked increase in circulating IgM autoantibodies and total serum IgM levels. Thus, CXCR4 expressing Treg cells control numbers of IgM autoantibody production by B1-B cells and the overall serum IgM levels.

## Results

### Phenotypic features and stability of CXCR4 expressing Treg cells

To enable quantification, tracking and “time stamping” of CXCR4-expressing Treg cells we generated *Cxcr4^CreERT2-GFPWT^Foxp3^Thy1.1^* mice, in which CXCR4^+^ Treg cells coexpressed Thy1.1 and GFP (Supplementary Figure 1a). In agreement with previous reports, Thy1.1^+^ Treg cells were highly enriched within total CD4 T cell population in the BM (Figure 1a) starting from 2 weeks of age (Supplementary Figure 1b) in contrast to other “Treg-rich” tissues such as skin, colon or adipose tissue with their markedly slower age-dependent buildup of Treg numbers. This result was confirmed by analyzing *Cxcr4^WT/WT^Foxp3^Thy1.1^* mice harboring two functional copies of *Cxcr4* gene (Figure 1a). CXCR4^+^ Treg cells were most represented within BM, skin and peripheral blood Treg populations (Figure 1b). Express labeling of intravascular CD45^+^ cells upon intravenous administration of a fluorophore-conjugated CD45 antibody suggested that Treg cells were residing primarily in the BM parenchyma rather than vasculature in contrast to other heavily vascularized organs such as lung and liver (Figure 1c). Nevertheless, both BM and splenic Treg cells were similarly replenished from hematogenous sources as revealed by parabiosis of CD45.1 and CD45.2 *Foxp3^DTR-GFP^* mouse pairs analyzed at 2 and 4 weeks post-surgery (33% and 38% “mixing”, respectively), in contrast to markedly more pronounced tissue residency exhibited by colonic Treg cells (Figure 1d; Supplementary Figures 1d,e).

**Figure 1.**
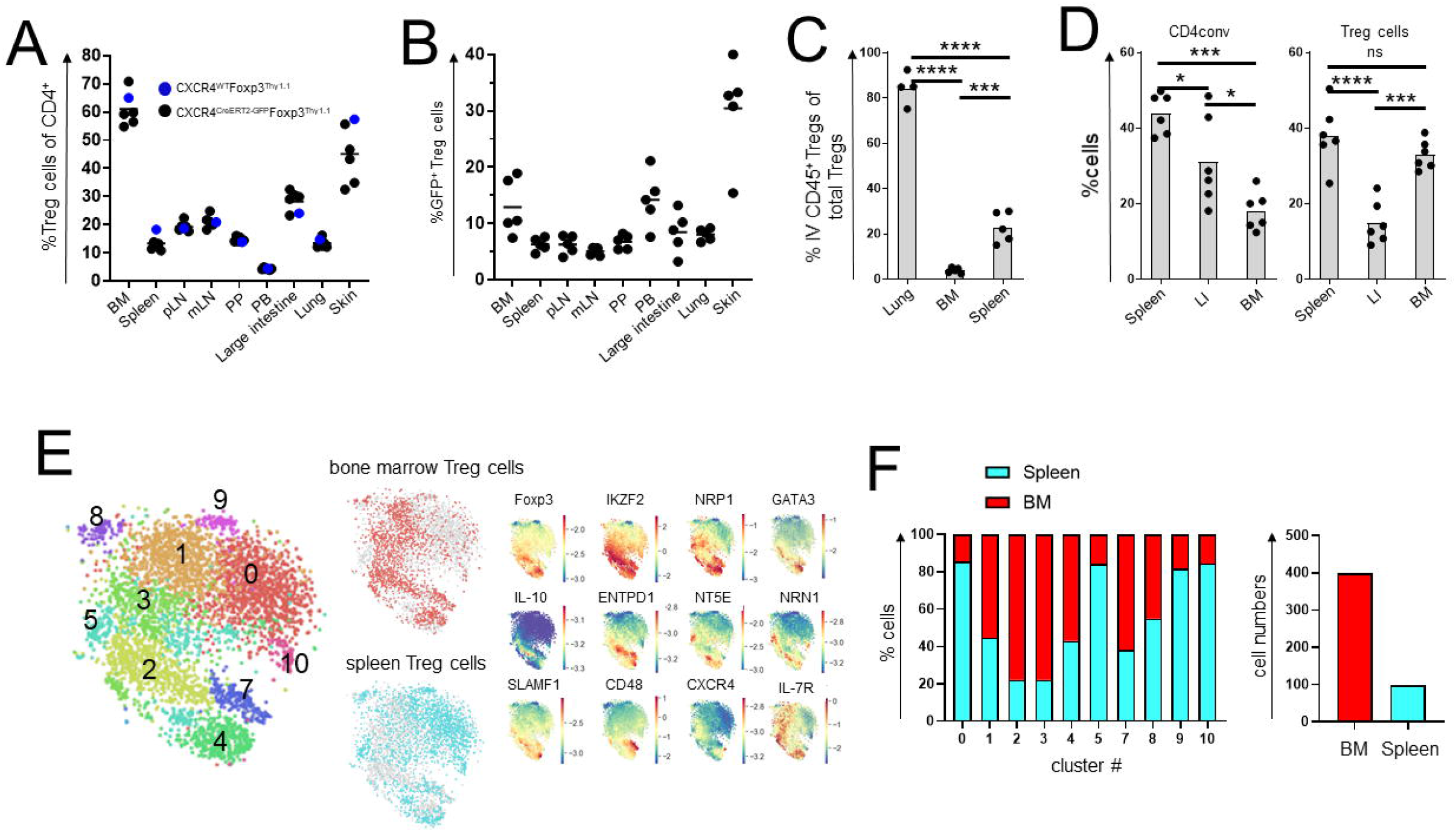
Features of CXCR4^+^ BM Treg cells. Distribution of total (A) and CXCR4^+^ Treg cells (B) in different tissues. pLN – peripheral lymph nodes; mLN – mesenteric lymph nodes; PP - Peyer’s patches; PB – peripheral blood. C) Analysis of intra-(IV^+^) and extravascular Treg cells in *Foxp3D^TR-EGFP^* mice labeled with CD45-BV570 for three min in the indicated tissues. \*\*\**p*< 0.001; ANOVA with multiple comparisons. D) Analysis of CD4conv and Treg cells in CD45.1^+^ and CD45.2^+^ *Foxp3D^TR-EGFP^* parabionts two weeks after surgery. LI - large intestine; BM - bone marrow. E) t-SNE visualization of scRNA- seq of spleen and BM Treg cells. Clustering analysis (left) and select gene distribution (right). F) Distribution of BM and spleen Treg cells among scRNA-seq clusters (left) and CXCR4^+^ Treg numbers (right). The data from one representative experiment of two independent experiments are shown in A-D.

For in-depth phenotyping we performed scRNA-seq analysis of BM and splenic Treg cells and their conventional counterparts (CD4conv) which showed relative enrichment of BM Treg cells in clusters 2 and 3 and their underrepresentation in cluster 0 in comparison to splenic Treg cells (Figure 1e,f; Table 1). While overall CXCR4 transcript levels were low, they were notably increased among BM Treg cells and correlated with increased expression of *Ikzf2* (Helios), *Ifngr1, Il10, Entpd1* (CD39), *Nt5e* (CD73), *Nrn1* (neuritin), *Nrp1* (neuropilin-1), *Tnfrsf18* (GITR), *Icos, Hif1a,* and *Gata3* genes (Figure 1e,f; Table 2). Thus, in BM Treg cells CXCR4 expression is associated with increased expression of suppressor function related genes.

### Induced CXCR4 ablation in Treg cells results in their selective BM depletion

To explore the functional significance of CXCR4 expression in Treg cells we generated *Foxp3^CreERT2-GFP^Cxcr4^FL/F^* mice, in which CXCR4 ablation in Treg cells can be induced upon tamoxifen administration, and control *Foxp3^CreERT2-GFP^Cxcr4^WT/WT^* mice hereafter termed FL and WT, respectively. Analysis of adult FL mice fed tamoxifen diet for 4-5 weeks showed a pronounced reduction in BM Treg cells (Figure 2a). While germ-line deficiency in a single *Cxcr4* allele did not have a measurable effect on Treg numbers in the BM and elsewhere (Figure 1a), its tamoxifen-induced acute ablation in Treg cells resulted in their moderate but statistically significant decrease in the BM (Figure 2a). Therefore, FL and WT mice were used for all subsequent studies. The observed impact of the induced loss of CXCR4 on Treg numbers in the former was highly selective to the BM, as Treg cells at all other sites were unaffected (Figure 2b). It is noteworthy that despite high levels of CXCR4 expression by thymic Treg cells, a mild decline in their numbers was merely a trend not reaching statistical significance (Figure 2b; Supplementary Figure 2a,b) and CXCR4 loss did not have noticeable effects on thymic Treg progenitors (Supplementary Figure 2c).

**Figure 2.**
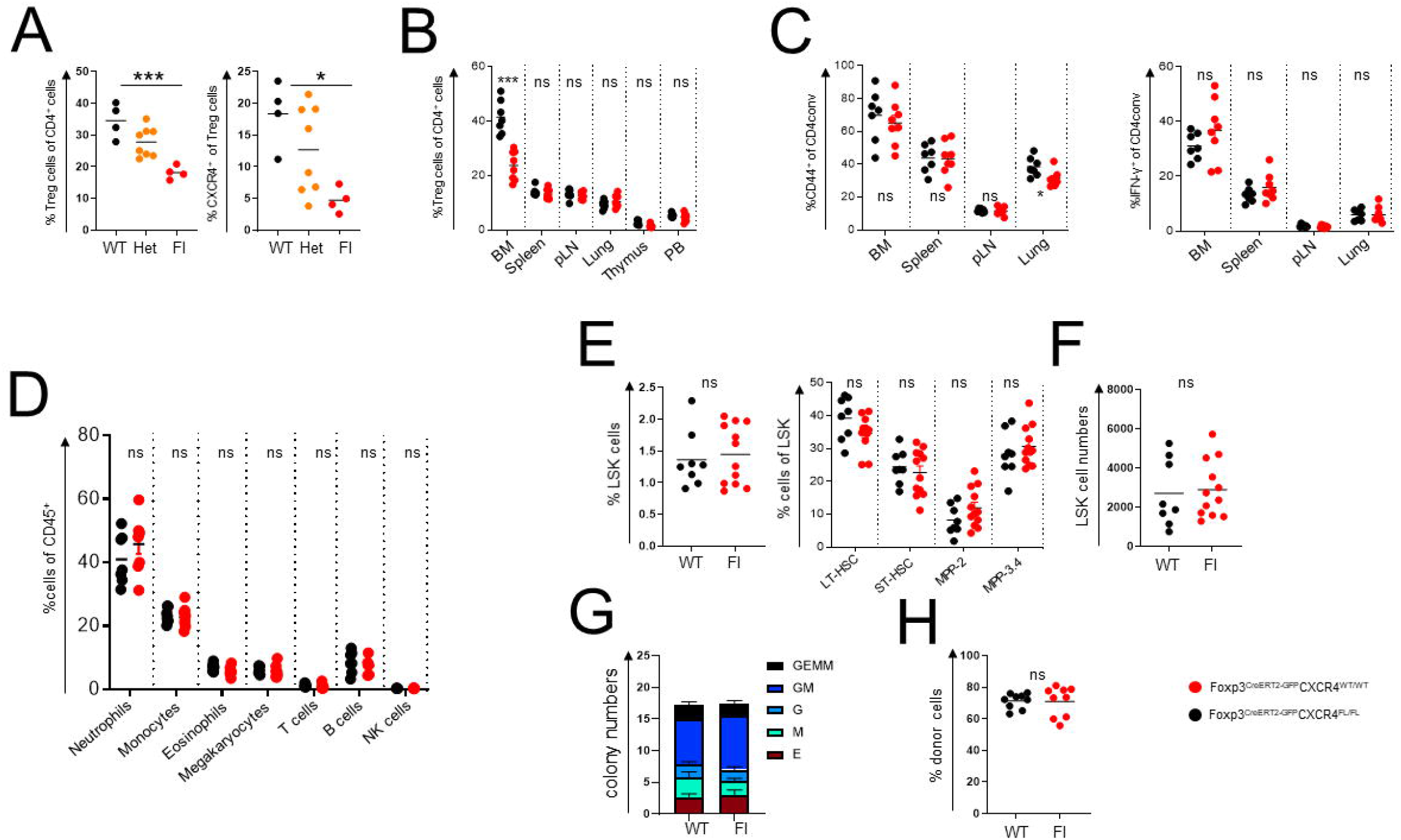
Induced deletion of CXCR4 in Treg cells. A, B) Analysis of total and CXCR4^+^ Treg subsets in the BM (A) and other tissues (B) of tamoxifen-treated WT and FL mice (one representative of two independent experiments is shown). C) Analysis of activated CD44^hi^CD62L^lQ^ (left) and IFNγ^+^ cells (right) in tamoxifen-treated WT vs FL mice (one representative of two independent experiments is shown). D-F) Analysis of hematopoietic lineages (D) and Lin^-^Sca-1^+^c-Kit^+^ (LSK) cells (E,F) in tamoxifen-treated WT and FL mice (one representative of four independent experiments is shown). LT – long term; ST – short term; HSC – haemopoietic stem cells; MPP - multipotent progenitors. G) *In vitro* colony formation by BM cells from WT and FL mice (n=6 of each genototype) (one representative of two independent experiments is shown). H) Analysis of competitive HSPC engraftment. CD45.2^+^ BM cells from tamoxifen-treated WT or FL and CD45.1^+^ BM cells were transferred at a 1:1 ratio into lethally irradiated CD45.1^+^ recipients and peripheral blood was analyzed 12 weeks later (one representative of two independent experiments is shown); ns – not significant, \**p*<0.05, Student’s *t* test.

The induced loss of CXCR4 in Treg cells did not result in any measurable generalized defect in their function as tamoxifen-treated FL mice remained clinically healthy with no significant changes in CD4 and CD8 T cell activation, B cell subset composition or pro-inflammatory cytokine production in the secondary lymphoid tissues and non-lymphoid organs such as lung (Figure 2c; Supplementary Figure 2e,f; data not shown).

While cell surface phenotype and proliferative activity of Treg cells were unchanged, we noted a mild increase in CD69 expression limited to the BM Treg subset – a likely reflection of compensatory activation caused by their diminished numbers (Supplementary Figure 2f).

Next, we sought to assess whether the loss of CXCR4 expression by Treg cells and their consequent decrease in the BM affected generation and numbers of innate immune cell lineages. We found major leukocyte subsets numerically unchanged in tamoxifen-treated FL mice (Figure 2d) with no detectable extramedullary hematopoiesis (Supplementary Figure 2d). Contrary to a previous suggestion that BM Treg cells control turnover and numbers of HSCs in a CXCR4-dependent manner (Hirata *et al.*, 2018), we found Lin^-^Sca1^+^cKit^-^ (LSK) cells and their subsets, including long-term HSCs defined as CD150^+^CD48^-^ LSK cells were unaffected by the induced Treg-restricted CXCR4 deficiency (Figure 2e,f). Furthermore, *in vitro* colony-forming activity of HSCs isolated from tamoxifen-treated WT and FL mice (Figure 2g) and their potential for competitive reconstitution of lethally irradiated hosts, when admixed with CD45.1^+^ HSCs at a 1:1 ratio, were comparable (Figure 2h).

### CXCR4 loss by Treg cells selectively affects the BM B cell compartment

Upon further characterization of the immune status of mice harboring CXCR4-deficient Treg cells we observed a notable and highly selective increase in serum IgM while amounts of other Ig isotypes including IgG1, IgG2b and IgG2c were comparable in tamoxifen-treated FL vs. WT mice (Figure 3a). Consistently, sera of FL mice prior and post tamoxifen treatment also showed a reproducible rise in IgM levels (Figure 3b). To test whether the latter reflected numerical changes in a specific B cell subset(s) despite largely unaltered bulk peripheral and BM B cell population sizes, we performed a finer grain flow cytometric analysis of immature and mature B cells in WT and FL mice (Figure 3c and Supplementary Figure 3b). We observed a significant reduction in the fraction, but not the numbers of pre-B cells and an increase in both proportion and absolute numbers of mature B220^+^CD43^-^IgM^+^IgD^+^B cells in the BM (Figure 3c). In contrast, tamoxifen treated FL and WT harbored comparable splenic B cell subsets including transitional cells, follicular and marginal zone B cells, and total B cell populations in the lymph nodes, spleen and lung as noted above (Supplementary Figure 2e; Figure 3d). Thus, deletion of CXCR4 in Treg cells led to an increase of mature B cells in the BM and increased serum IgM levels.

**Figure 3.**
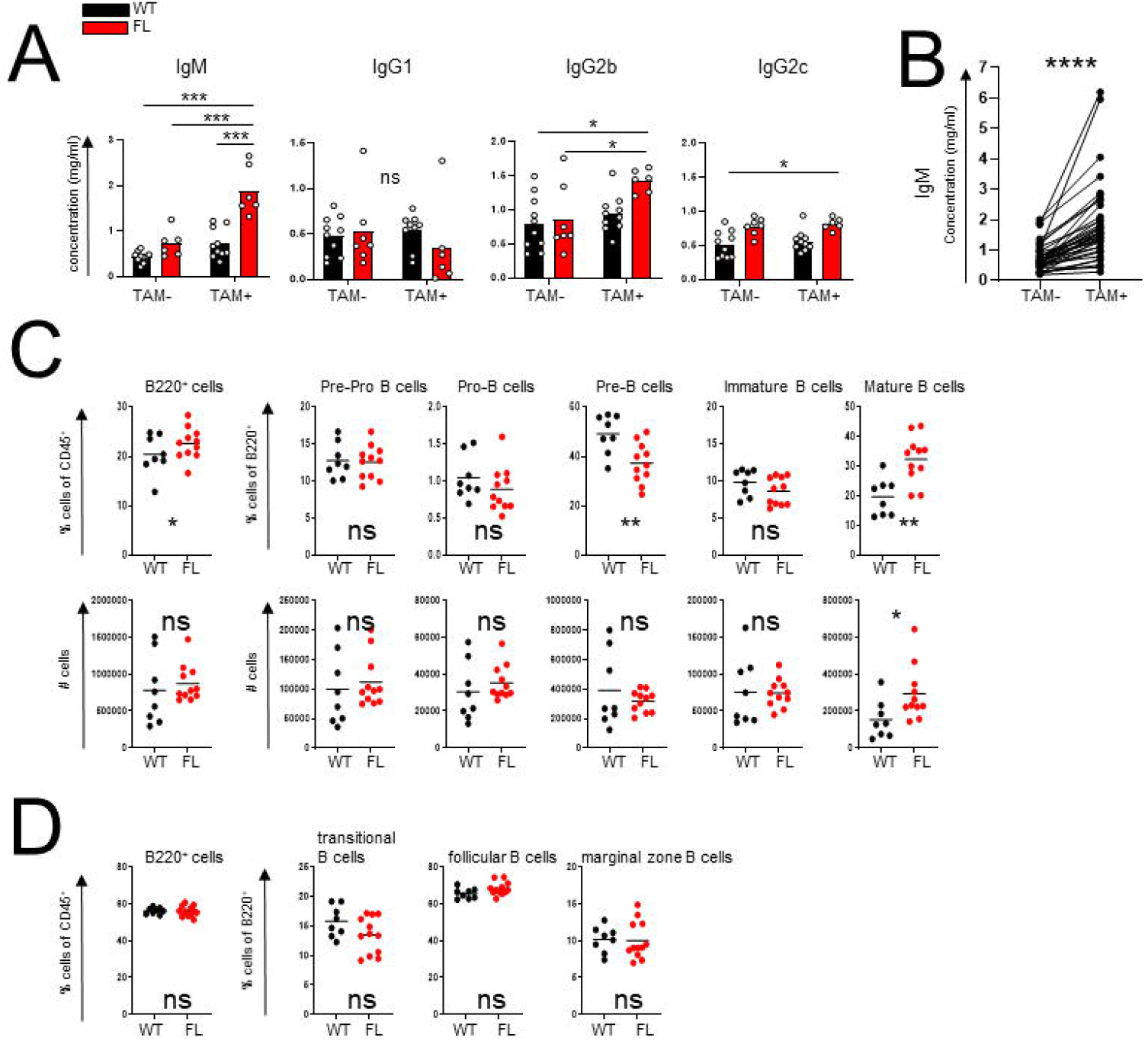
CXCR4 deletion in Treg cells results in an increase in serum IgM and mature BM B cells. A) Serum Ig isotype levels of WT and FL mice before and after tamoxifen induced CXCR4 deletion (one representative experiment out of four experiments). B) Aggregate changes in IgM levels in FL mice before and after tamoxifen administration (four independent experiments). C) BM B cell subsets shown as fraction of CD45^+^ or B220^+^ cells (upper panels) and absolute numbers (lower panels) (one representative experiment out of five). (D) Splenic B cell subsets in WT and FL mice (one representative experiment out of three). ns, not significant; * *P* < 0.05, ** *P* < 0.01 and *** *P* < 0.001. ANOVA with multiple comparisons (A); paired Student’s t-test (B); unpaired Student’s *t* test (C,D).

### CXCR4 expressing Treg cells control B-1 cells in the bone marrow

To identify the potential source of the increased IgM amounts in FL mice, we performed ELISPOT assay of BM and spleen cells (Figure 4a). We noted a significant and highly selective increase in IgM-, but not IgG-producing antibody secreting cells (ASCs) in the FL vs WT BM (Figure 4a) while the frequencies of both IgM and IgG-producing ASCs in the spleen were comparable WT and FL mice (Figure 4b). These observations suggested that higher levels of serum IgM we observed in FL mice resulted from increased ASCs in the BM of FL compared to WT mice.

**Figure 4.**
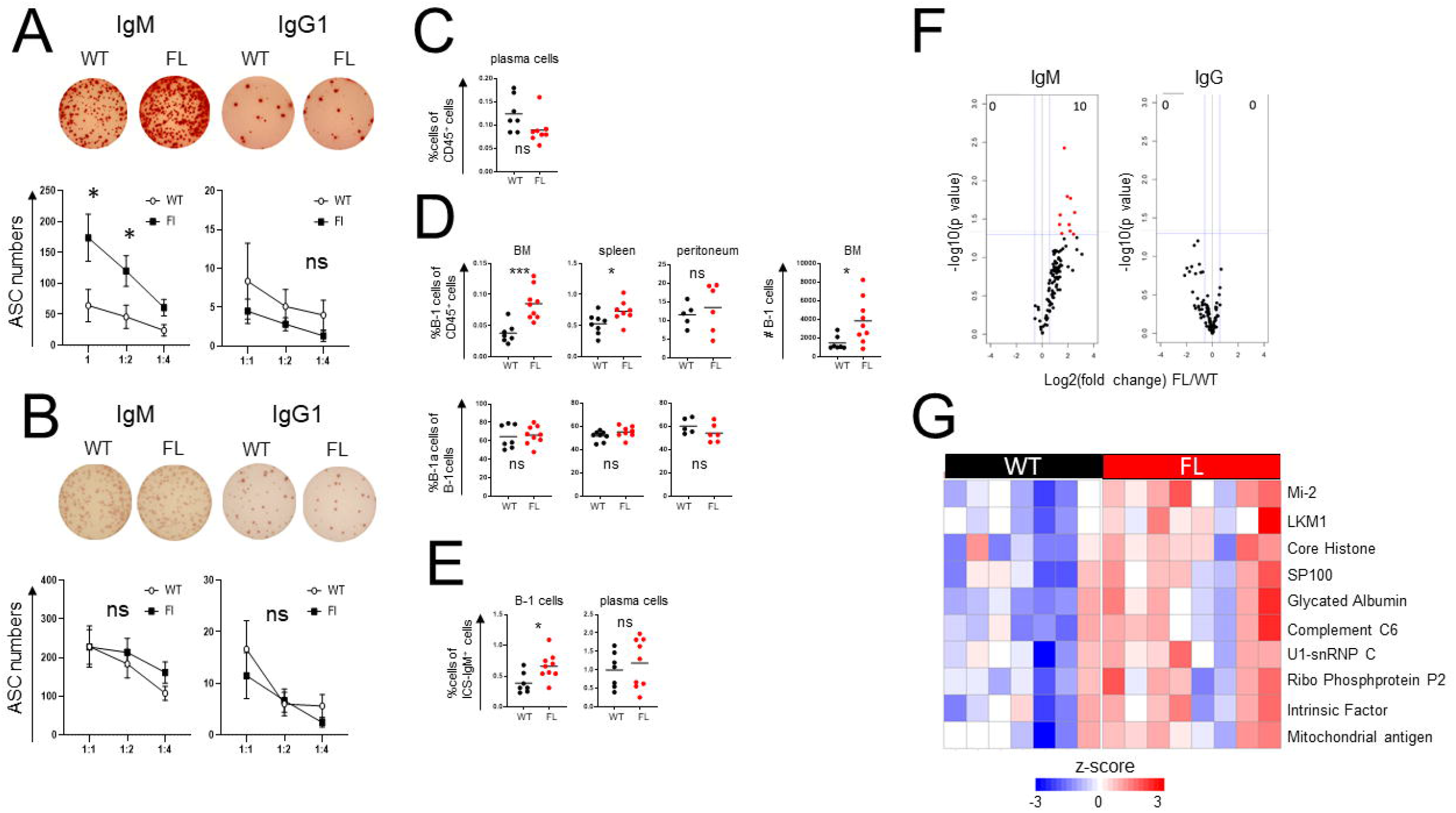
BM Treg cells control B-1 cells. ELISPOT analysis of BM (A) and spleen cells (B) of tamoxifen-treated WT and FL mice. Quantification of plasma cells in the BM (C) and B-1 cells in the indicated tissues (D) in tamoxifen-treated WT and FL mice. ACS – antibody-secreting cells. (E) Flow cytometric analysis of intracellular IgM expression. The percentages of intracellular IgM^+^ cells within indicated cell subsets are shown. (F,G) Analysis of reactivity of serum antibodies in tamoxifen-treated WT and FL mice with a 128-autoantigen microarray. Volcano plots (F, left: IgM; right: IgG; red: highly ranked) and heat map (G) of the mean intensity of autoantibody reactivity in FL vs. WT mice. For A-E, one representative experiment is shown out of at least two. ns – not significant; \**P* < 0.05, *** *P* < 0.001, Student’s *t* test.

Since plasma cells and B-1 B cells are thought to serve as major IgM producing cell types in unchallenged mice, we assessed their frequencies in WT and FL mice (Figure 4c). While we observed comparable numbers of CD138^+^B220^-^ plasma cells (Figure 4c), we found markedly higher proportion and numbers of B220^-^CD19^+^CD23^-^CD43^+^ B-1 cells in the BM, but at best marginal increase in the spleen and no change in the peritoneal cavity (Figure 4d; Supplementary Figure 4a). Consistent with the possibility of B-1 cells contributing to heightened serum IgM levels, proportion of B-1 cells, but not plasma cells expressing intracellular IgM was also increased in tamoxifen-treated FL vs WT mice (Figure 4e). We have not observed increases in BM B-1 cells or mature B-2 cells upon transient depletion of Treg cells in *Foxp3^DTR^* mice (Supplementary Figure 4b).

Considering a known bias of B-1 cell specificity towards “self”, we analyzed reactivity of circulating IgM and IgG antibodies in these mice using a panel of 128 known autoantigens. While the titers of IgG1 autoantibodies were comparable in tamoxifen-treated FL vs WT mice, in the former we observed increased amounts of IgM autoantibodies directed against a range of autoantigens; most prominent among those were Mi-2, LKM1, core histone and SP100 (Figure 4f, g; Supplementary Figure 4c). These results were confirmed by ELISA with the most frequent autoantigens in this panel (data not shown). Furthermore, ELISPOT assay revealed increased frequency of BM B-1 cells, producing autoantibody against a pool of these predominant autoantigens, in FL vs WT mice while no significant autoantibody production by BM B-2 cells was observed (Supplementary Figure 4d). These results suggest that under physiologic conditions BM Treg cells regulate, in a CXCR4 dependent manner, the levels of self-reactive serum IgM through control of their production by B-1-B cells in the BM.

## Discussion

Besides exerting their widely recognized generic suppressor activity, Treg cells residing in lymphoid and non-lymphoid tissues also contribute to certain specific aspects of tissue functions in physiological and pathological settings (Munoz-Rojas and Mathis, 2021; Panduro et al., 2016). These specialized organ-specific “homeostatic” roles of Treg cells remain poorly understood in part because the majority of studies have relied on wholesale depletion or constitutive genetic perturbations of Treg cells. Pronounced systemic autoimmunity and inflammation caused by the former can obscure more nuanced functions of Treg cells while the latter may affect the “baseline” state of developing tissue. Here, we show that temporally controlled interference with the migration of Treg cells to, and their residence within a given tissue can alleviate these confounders and reveal novel, previously unappreciated facets of Treg biology. Through inducible ablation of the chemokine receptor CXCR4 in Treg cells causing their partial and highly selective depletion in the BM, we uncovered a novel role of CXCR4-expressing BM Treg cells in controlling B-1 cell numbers and autoantigen-specific IgM production by these cells exclusively in the BM. This is consistent with the notion that BM B-1 cells serve as major IgM producers and main contributors to circulating IgM pool at physiologic conditions (Choi et al., 2012; Reynolds et al., 2015; Savage et al., 2017). In agreement with the aforementioned notion of “masking” effects of systemic Treg depletion, the latter failed to increase BM B-1 or mature B-2 cell numbers.

Importantly, the observed effects of CXCR4 were not associated with any noticeable changes in immune balance outside the BM, in numbers of HSCs or their colony forming and BM reconstitution capacity. These findings differ from previous studies, which suggested that a sizeable proportion of BM Treg cells are found in the proximity of the endosteal surface – a site of primitive HSC niches, and implicated IL-10 and adenosine production by BM Treg cells in HSC maintenance (Fujisaki et al., 2011; Hirata *et al.*, 2018). Depletion of Treg cells induced by DT in *Foxp3^DTR^* mice or by depleting CD25 antibodies has been associated with a reduction in the numbers of allo-HSCs and progenitor cells (HSPC) (Fujisaki *et al.*, 2011). Furthermore, constitutive deletion of a *Cxcr4* conditional allele in Treg cells was shown to increase BM cellularity, HSPC and HSC numbers, and enhance BM reconstitution upon BM transplantation (Hirata *et al.*, 2018). Beyond aforementioned potential effects of pronounced inflammatory responses induced upon Treg ablation on HSCs, additional explanation for the apparent discrepancy between these previous findings and our observation that CXCR4-expressing Treg cells were dispensable for HSC maintenance is potential developmental alterations in the BM of mice harboring CXCR4-deficient Treg cells. In support of this possibility, we observed pronounced enrichment of Treg cells among overall CD4 T cell population in the BM as early as 2 weeks of life as opposed to their much slower build-up in other “Treg-rich” organs.

While the sparsity of both B-1 cells and Treg cells in the BM presents a major obstacle to a rigorous assessment of their relative proximity, we propose that BM Treg and B-1 cells localize to the same niche guided by the their shared expression of CXCR4 with CXCL12 expressing stromal cells serving as their meeting points in the BM. CXCR4 is known to be expressed by all subsets of B cells thought B cell ontogeny and mice deficient in CXCR4 or CXCL12 display defects in B cell development (Epling-Burnette et al., 2007; Tachibana et al., 1998; Zou et al., 1998). Importantly, selective loss of CXCR4 in B cells results in premature migration of B cell precursors from the BM and their localization in splenic follicles (Nie *et al.*, 2004). It has also been shown that the homing of mature B cells to the BM depends on CXCR4 expression (Beck et al., 2014; Nie *et al.*, 2004; Pereira et al., 2009). Although these findings provide rationale for direct interactions between BM Treg cells and B cells, their indirect interactions through stromal cells are also possible considering a role for stromal cells in B cell development (Wu et al., 2008). Although the identity of a Treg derived factor or factors acting directly or indirectly upon B-1 cells remains unknown, we found the two major “suspects” – neuritin recently implicated in control of GC B cells by follicular Treg cells (Gonzalez-Figueroa et al., 2021) and Treg-derived IL-10 – are fully dispensable for Treg mediated control of serum IgM levels (data not shown).

Despite considerable research efforts, control mechanism of B-1 B cells and natural IgM levels remained unknown (Baumgarth, 2017). Our studies demonstrate that Treg cells, in a CXCR4 dependent manner, control IgM autoantibody production by B-1 B cells highlighting a novel mechanism of cell-extrinsic regulation of BM B-1 B cells and physiologic amounts of serum IgM and their self-reactivity. These results may have implications for understanding the pathogenesis of autoimmune and B cell lymphoproliferative diseases such as chronic lymphoproliferative leukemia (CLL), which has been linked to a human counterpart of mouse B-1 B cells (Hayakawa et al., 2016; Herve et al., 2005).

## Methods

### Mice

*Foxp3^CreERT2^, Foxp3^Thy1.1^, Foxp3^DTR^* and *CXCR4^CreERT2^* mice were described elsewhere (Kim et al., 2007; Liston et al., 2008; Rubtsov et al., 2010; Werner et al., 2020). *CXCR4*^Fl^, B6 CD45.1 mice were purchased from the Jackson Laboratory. The experimental mice on C57BL/6J (B6) background were maintained in the specific pathogen-free animal facility at Memorial Sloan Kettering Cancer. All animal experiments were approved by institutional animal care and use committee. 5-12 wk-old mice were used for all experiments unless mentioned otherwise and were age and sex matched in individual experiments. For induced of CXCR4 deletion in Treg cells, mice were placed on a tamoxifen citrate–containing diet (TD.130860; Envigo) for 4-5 wks. For diphtheria toxin induced Treg ablation, *Foxp3*^DTR^ mice received i.p. injection of 500 ng DT (List Labs) on days 0 and 1 and tissues analyzed at day 10.

### Parabiosis and IV labeling

Female CD45.1 and CD45.2 B6 mice underwent surgery to establish parabiosis, as previously described (Fan et al., 2018; Kamran et al., 2013). For intravenous (IV) labeling of CD45^+^ cells, mice received IV injection of 3 μg of a Brilliant Violet 570-conjugated CD45 antibody (BioLegend) and sacrificed 3 min later and the fraction of IV-labeled cells was quantified.

### Tissue preparation and cell isolation

Spleen, lymph nodes, and thymi were mechanically dissociated with the back of a syringe plunger and filtered through a 100-μm cell strainer (Corning). For isolation of colon lamina propria cells, the colon was first incubated with EDTA solution containing PBS, 2% FBS, 10mM HEPES buffer, 1% penicillin-streptomycin, 1% L-glutamine, 1mM EDTA and 1mM DTT at 37°C for 15 min while shaking. After centrifugation, the intraepithelial fraction was removed. Colon tissue samples devoid of intra-epithelial cells as well as lung and skin samples were enzymatically digested with 1 mg/ml Collagenase A from *Clostridium histolyticum* (Roche) and 1 unit/ml DNaseI (Sigma-Aldrich) with 2% FBS, 1% penicillin/streptomycin, 1% L-glutamine, and 10 mM HEPES in the presence of 0.25-inch ceramic spheres (MP Biomedicals) for 45 minutes at 37°C while shaking. Digested tissue was filtered through 100-μm cell strainers and colon and lung cells were further enriched by 40% Percoll (GE Healthcare) centrifugation. For isolation of BM cells, one tibia and one femur from each mouse were cut at their ends and placed into a 0.2 ml microcentrifuge tube which was previously pierced by an 18G needle (Amend et al., 2016). The microcentrifuge tube was placed into a 1.5 ml microcentrifuge tube and centrifuged at 2,000 X g for 1 minute.

Spleen, BM, lung and liver cell suspension were subjected to red blood cell (RBC) lysis before down-stream applications.

### Flow cytometric analysis and cell sorting

Single-cell suspensions were re-suspended in PBS and stained with Ghost Dye Violet 510 (Tonbo Biosciences) for dead-cell exclusion in the presence of anti-CD16/32 (Tonbo Biosciences) to block binding to Fc receptors for 10 min at 4°C. Cell-surface antigens were stained for 20 min at 4°C in FACS buffer (PBS, 2% BSA, 2 mM EDTA, 1% L-Glutamine and 10 mM HEPES). Cells were analyzed unfixed or fixed with 1% paraformaldehyde (Electron Microscopy Sciences) for later analysis or fixed and permeabilized with the BD Cytofix/Cytoperm kit for intracellular staining. Fixation, permeabilization, and intracellular staining were performed as recommended by the manufacturer. All cells were re-suspended in FACS buffer and filtered through 100-μm nylon mesh before analysis on the flow cytometer. To aid acquisition, samples were treated with 40 U/mL DNase I for 10 min at 37°C before acquisition. 123count eBeads (eBioscience) were added at 5,000 beads per sample to quantify absolute cell numbers. All flow cytometry data were collected on a BD LSRII (Becton Dickinson) flow cytometer or the Aurora spectral analyzer (Cytek) and analyzed using FlowJo 10 software (TreeStar Software). Cell suspensions were stained with fluorescence-tagged antibodies as described in STAR methods. To assess cytokine production after ex vivo re-stimulation, single-cell suspensions were incubated in round-bottom 96-well plates for 3 h at 37°C with 5% CO2 in the presence of 50 ng/ml PMA and 500 ng/ml ionomycin with 1 μg/ml brefeldin A (all Sigma-Aldrich) and 2 μM monensin (BioLegend). For intracellular staining of IgM, cells were first incubated with an unconjugated IgM antibody (10μg/ml; clone II/41, eBioscience) together with fluorophore-tagged panel of surface marker-specific antibodies for 30 min on ice. Then the cells were fixed and permeabilized (BD Cytofix/Cytoperm kit) and stained intra-cellularly with PE conjugated anti-IgM antibody (clone II/41, eBioscience). For sorting CD4+ cells (CD4conv and Treg cells), cells were first enriched using Dynabeads FlowComp mouse CD4 (Invitrogen), then stained with fluorochrome-conjugated CD45, TCRβ, CD4 and GR1 antibodies and sorted using a BD FACSAria II cell sorted (Treg cells were identified as GFP+CD4+ cells).

### Serum Ig ELISA

ELISA was performed for quantification of the concentration of total serum Ig. For ELISA experiments mouse peripheral blood collected by retroorbital bleeding under isoflurane anesthesia or inferior vena cava (IVC) blood collected by terminal bleeding was collected into BD SST microcontainer tubes and sera were harvested after centrifugation. Nunc-Immuno MaxiSorp F-bottom 96-well plates were coated with 50 μL/well capture antibodies in PBS pH 7.4 overnight at 4°C. The plates were then washed 3 times with PBS containing 0.05% Tween-20 (Sigma-Aldrich), blocked with 150 mL ELISA/ELISPOT Diluent (Life Technologies Corporation) for 3 hrs, and washed with PBS containing 0.05% Tween-20. 50 μL of sera at appropriate dilutions was added and incubated overnight at 4°C. The plates were then washed with 0.05% PBS-Tween and incubated with 50 μL of Goat Anti-Mouse Ig-HRP at room temperature for 2 hrs, washed 7 times with 0.05% PBS-Tween and developed with 50 μL of TMB solution (SeraCare, 5120-0047) at room temperature. The colorimetric reaction was stopped with 50 μL of 1M H_3_PO_4_ (Sigma-Aldrich, P5811) after 1-5 minutes of adding TMB and absorbance at 450 nm was measured with a Synergy HTX plate reader (BioTek). Concentrations of antigens were determined using standard curves constructed with purified recombinant proteins and calculated with Gen5 3.02.2 (BioTek).

### Measurement of serum autoantibodies

Serum autoantibodies were analyzed using an autoantigen array chip containing 128 antigens and controls at Microarray and Immune Phenotyping Core Facility (UT Southwestern Medical Center). Briefly, the autoantigens and control proteins are printed in duplicates onto Nitrocellulose film slides (Grace Bio-Labs). 2ul serum samples were pretreated with DNAse-I and then diluted 1:50 in PBST buffer for autoantibody profiling. The diluted serum samples were incubated with the autoantigen arrays, and autoantibodies binding with antigens on arrays were measured with cy3-conjugated anti-human IgG (1:2000, Jackson ImmunoResearch Laboratories) and cy5-conjugated anti-human IgM (1:1000, Jackson ImmunoResearch Laboratories), using a Genepix 4200A scanner (Molecular Device) with laser wavelength of 532 nm and 635 nm. The resulting images were analyzed with Genepix Pro 7.0 software (Molecular Devices). The median of the signal intensity for each spot are calculated and subtracted the local background around the spot, and data obtained from duplicated spots were averaged. The background subtracted signal intensity of each antigen is normalized to the average intensity of the human IgG or IgM controls, which were spotted on the array as internal controls. Finally, the normalized fluorescence intensity (NFI) was generated as a quantitative measurement of the binding capacity of each antibody with the corresponding autoantigen. Signal-to-noise ratio (SNR) is another quantitative measurement of the true signal above background noise. SNR values equal to or greater than 3 were considered significantly higher than background, and therefore true signals. The autoantibody which has the SNR value less than 3 in more than 90% of samples was considered negative and excluded from further analysis. The NFI value of the autoantibodies were used for statistical analysis and the heatmaps were generated using Cluster and Treeview software (http://bonsai.hgc.jp/~mdehoon/software/cluster/software.htm). Statistical analysis was performed using R package to identify differential expressed antibodies between different study groups.

### Generation of bone marrow chimeras

For generation of mixed BM chimeras, BM cells were isolated from WT and FL (*Foxp3*^Cre–ERT2^*CXCR4*^WT/WT^ or *Foxp3*^Cre–ERT2^*CXCR4*^FL/FL^, respectively) mice which were maintained on tamoxifen diet for 4-5 wks and from CD45.1 mice (as described above) which were subjected to T cell depletion using Dynabeads FlowComp Mouse Pan T (CD90.2) Kit (ThermoFisher Scientific). BM cells from WT or FL mice were mixed with the BM of the competitor donor at 1:1 ratio and then transplanted by intravenous injection into lethally irradiated CD45.1 mice (900 rad). Each recipient mouse received a total of 1×10^6^ cells of the BM mixture.

### Single cell-RNA seq analysis

FACS purified cell subsets were labeled by incubation for 30 minutes at 4°C with TotalSeq anti-mouse Hashtag antibodies (BioLegend), then washed, re-suspended at 1.5×10^3^ cells/μL in PBS, 0.04% BSA and finally mixed with the other labeled cells.

#### scRNA-seq sample preprocessing

Individual samples were loaded on 10X Genomics Chromium System. Libraries were prepared following 10X Genomics protocols (Chromium Single Cell 3’ Reagent Kits v2 Chemistry) and sequenced on NovaSeq 6000 (Illumina S2 flow cell, paired-end). FASTQ files were processed using cellranger (https://support.10xgenomics.com/single-cell-gene-expression/software/pipelines/latest/what-is-cell-ranger) and reads were aligned to the mouse genome mm10 from ENSEMBL GRCm38. Cells that contained a percentage of mitochondrial transcripts >5% were filtered out, resulting in 2,159 splenic regulatory T cells and 1,887 bone marrow regulatory T cells that passed QC metrics, with a median of 3,63 molecules/cell (in log10 scale). Cells with total molecule counts of <1,000, determined by lower mode of distribution of total molecules per cell, were additionally filtered out to remove putative empty droplets. Genes that were expressed in more than 10 cells were retained for further analysis. The resulting count matrices from all samples were then combined, resulting in a final set of 4,046 cells x 11,362 genes, and normalized to median library size, where library size is defined as total molecule counts per cell. The normalized data are then log transformed as log (counts + 1) for downstream analysis.

#### PCA

For dimensionality reduction and visualization of data, genes with very low or very high expression of transcripts (log average expression <0.02 or >3 and a dispersion >0.15) were excluded. Principal component analysis was performed on the log-transformed normalized data using 20 principal components, where the number of principal components was determined by the knee-point in eigenvalues.

#### MAGIC imputation

To account for missing values in scRNA-seq due to the high dropout rate, the imputation was performed using MAGIC (van Dijk et al., 2018). The diffusion maps matrix was constructed using k = 30, ka = 10, and t = 3 as input parameters, where t specifies the number of times the affinity matrix is powered for diffusion.

#### Clustering and gene ranking

Clustering of cells was performed using PhenoGraph (Levine et al., 2015) and setting k = 40 nearest neighbors. A cluster containing cells with a low number of total molecules compared with other populations was removed. Significant differentially expressed genes in each cluster were identified using *t*-test implemented in Single-Cell Analysis in Python (SCANPY) (Wolf et al., 2018) with default parameters. For identification of Cxcr4 expressing cells in scRNA-seq datasets we used the MAGIC imputed matrix and set a cutoff of −3.05 where any cell with a higher value than that is considered *Cxcr4* expressing (Cxcr4^+^) (446 cells). The rest of the population is considered *Cxcr4* nonexpressing (Cxcr4^-^).

### Colony forming assay

Bone marrow colony forming assay was performed by plating 10^4^ or 5×10^4^ cells on methylcellulose (MethoCult M3434, Stem Cell Technologies). GEMM (Granulocyte/Erythrocyte/Macrophage/Megakaryocyte), GM (Granulocyte/Macrophage), G (Granulocyte), M (Macrophage), CFU-E (Erythroid) and BFU-E (Erythroid) colonies were counted 2 wks after seeding.

### Statistical analysis and data availability

Statistical analyses were performed using Prism software version 8 (GraphPad, San Diego, CA). A statistical test was considered significant when p<0.05. scRNA-seq data of spleen and BM CD4+ cells have been deposited to the Gene Expression Omnibus under accession no. XXX.

## Acknowledgments

We thank A. Bravo and J. Verter for help with animal husbandry, B. Dhillon, C. Ariyan and Z. Wang for help in experiments (all from MSKCC), C. Vinuesa (The Australian National University, Canberra, Australia) for assistance with the neuritin mice experiments and C. Zhu (University of Texas Southwestern Medical Center, TX) for analysis of the autoantibody panels. We thank all Rudensky lab members for fruitful discussions. S.E. is a Cancer Research Institute Irvington Fellow supported by the Cancer Research Institute.

---

**Supplementary Figure 1.**
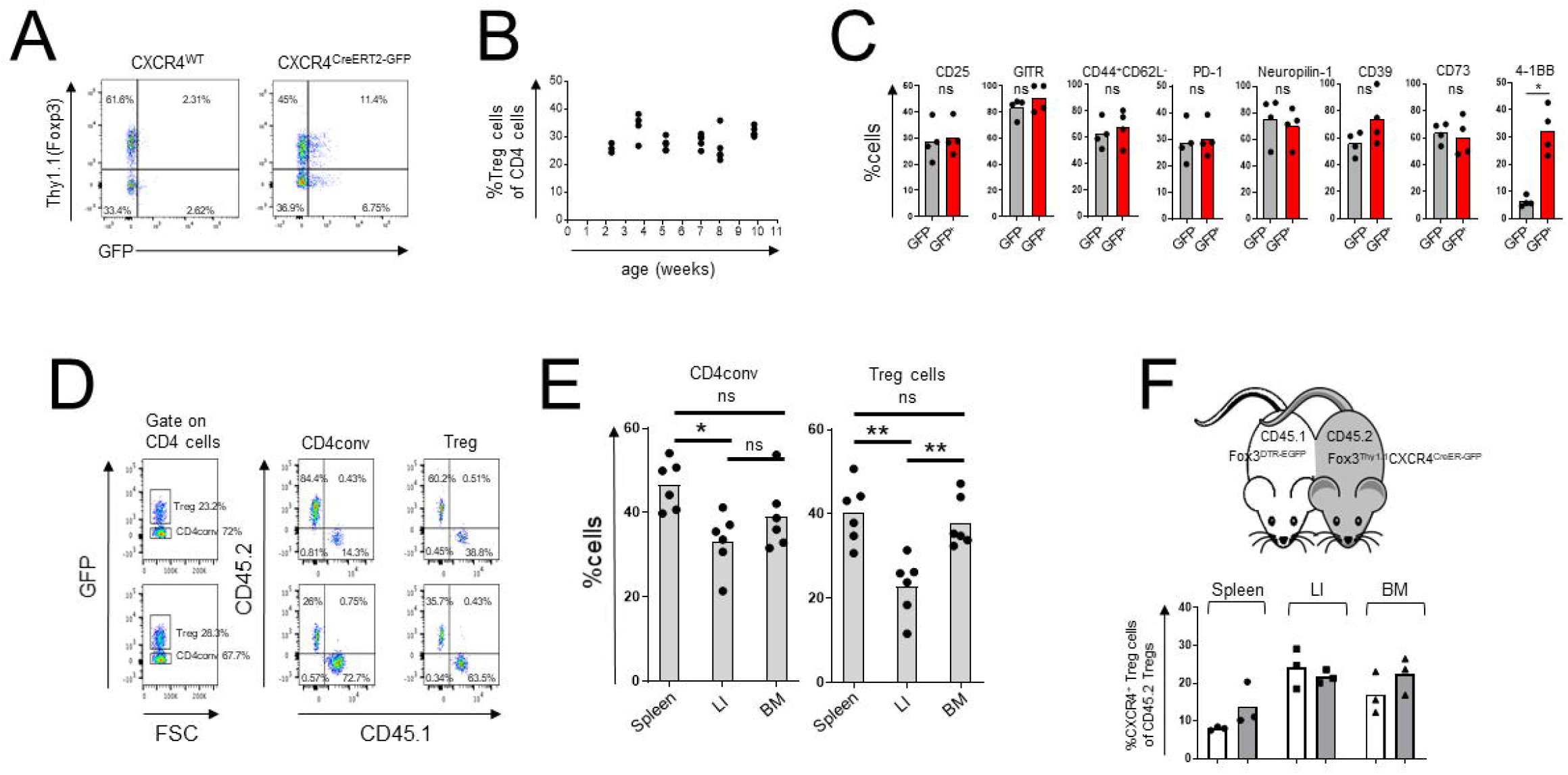
General properties of CXCR4-expressing BM Treg cells. A) Flow cytometric analysis of CXCR4^+^ Treg cells in *CXCR4C^reERT2-GFP^Foxp3^Thy1.1^* (right) and *CXCR4^WT^Foxp3^Thy1.1^* mice (left) after gating on CD4^+^TCRβ^+^ cells. B) Flow cytometric analysis of age-dependent abundance of Treg cells within CD4^+^ T cell population in *Foxp3*^DTR-EGFP^ mice. C) Flow cytometric analysis of expression of activation markers by CXCR4^+^ (Thy1.1^+^GFP^+^) and CXCR4^-^ Treg cells in *Cxcr4^CreERT2-GFP^Foxp3^Thy1.1^* mice. The data from one experiment representative of three independent experiments are shown. ns – not significant; \**p*<0.05, Student’s *t* test. D-F) Analysis of BM Treg cells in parabiotic mice. D) Gating strategies for flow cytometric analysis shown in Figure 1D. E) Flow cytometric analysis of Treg cells in CD45.1^+^ and CD45.2^+^ *Foxp3D^TR-EGFP^* parabionts four weeks after surgery. F) Analysis of Treg cells in Fox3^DTR-EGFP^CD45.1 and Fox3^Thy1.1^CXCR4^CreER-GFP^CD45.2 parabionts two weeks after surgery. The percent of CXCR4^+^ Treg cells out of CD45.2^+^ Treg cells was quantified in CD45.1 ^+^ (white bars) and CD45.2^+^ parabionts (grey bars). LI – large intestine.

**Supplementary Figure 2.**
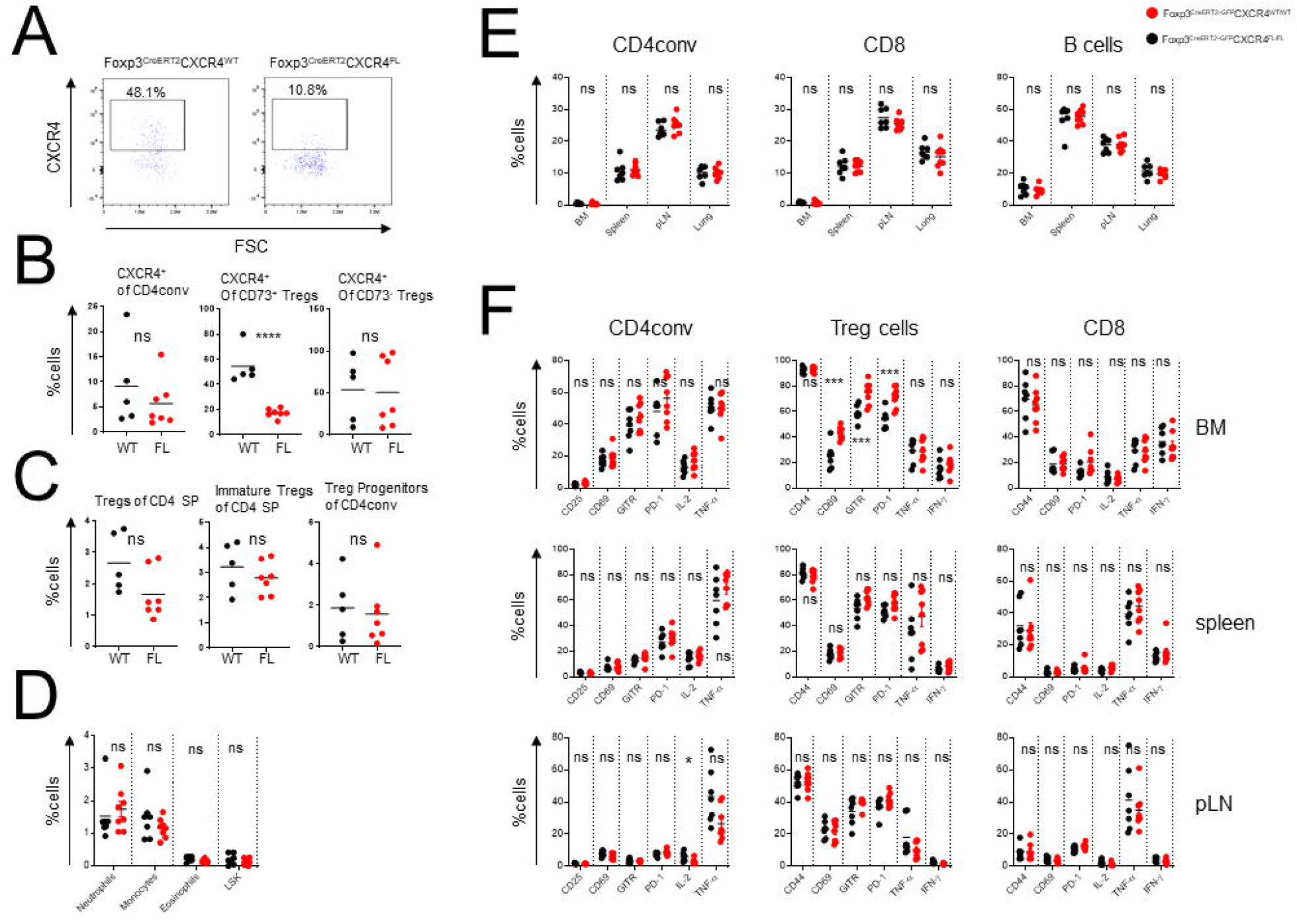
Thymic Treg cells and activation markers in FL mice. AC) Flow cytometric analysis of thymocytes in tamoxifen-treated WT and FL mice. CXCR4 expression by thymic Treg cells (A); proportion of CXCR4^+^ thymic Treg cells and CD4conv cells (B); frequencies of mature (Dump^-^CD4^+^TCRβ^+^CD8^-^CD73^-^ CD25^+^Foxp3^+^) and immature Treg cells (Dump^-^CD4^+^TCRβ^+^CD8^-^CD25^-^Foxp3^+^) and Treg progenitors (Dump^-^CD4^+^TCRβ^+^CD8^-^GITR^+^CD122^+^CD25^+^Foxp3^-^) (C). The dump gate includes staining for NK1.1, CD11b, CD19, MHC-II, TCRγδ and CD1d tetramer. Flow cytometric analysis of splenocytes (D), proportion of CD4conv, CD8 and B cells subsets in indicated tissues (E), expression of activation markers and cytokine production by CD4conv, Treg cells and CD8 cells in different tissues (F) in tamoxifen-treated FL and WT mice. Two right columns in each panel show intracellular cytokine production after *in vitro* stimulation. The data are shown for one representative of two (A-C) or three (D-F) independent experiments. sp – single positive. ns – not significant, \**p*<0.05, \*\*\**p*<0.001, Student’s *t* test.

**Supplementary Figure 3.**
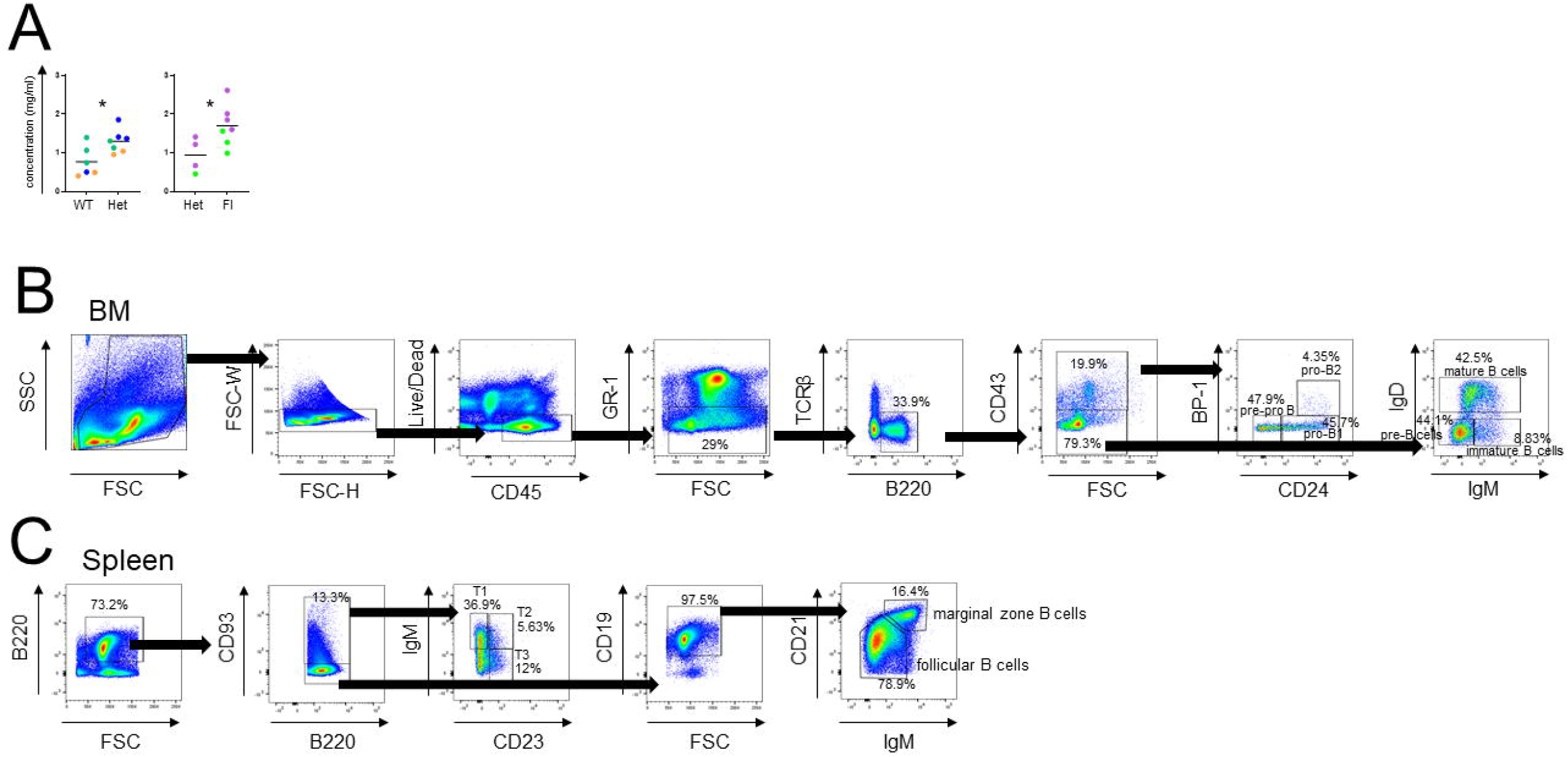
(A) Basal serum IgM levels of WT, *Foxp3^CreERT2^CXCR4^WT/FL^* (Het) and FL littermates (littermates are color coded). (B, C) Gating strategy for BM (B) and splenic (C) B cell subsets by flow cytometric analysis. \**p*<0.05, Student’s *t* test.

**Supplementary Figure 4.**
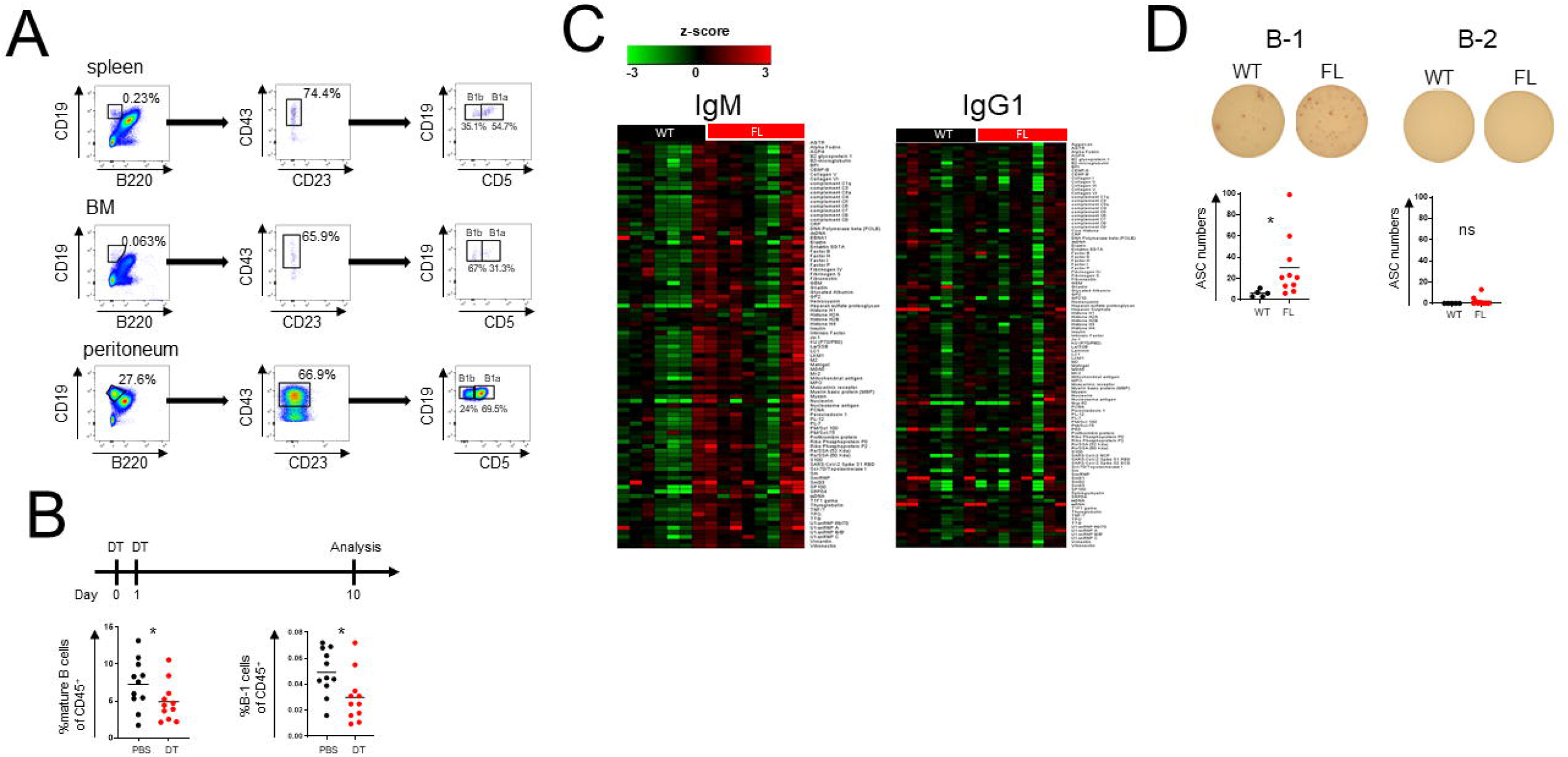
BM Treg cells regulate B-1 cells. A) B-1 cells gating strategy. B) Staining of mature B cells (left) and B-1 cells (right) in the BM in Fox3^DTR^ mice 10 days after two DT or PBS (control) injections. Data are combined from two independent experiments with at least five mice per group. C) Heatmap of IgM and IgG1 autoantibodies. D) ELISPOT analysis of BM B cells. BM B-1 (left) and B-2 cells (right) were sorted after CD19 magnetic bead enrichment and plated on ELISPOT plates precoated with five auto-antigens: core histone, ribo phosphoprotein P2, intrinsic factor, mitochondrial antibody and SP100. ACS – antibody-secreting cells. * *P* < 0.05, Student’s *t* test.

**Supplementary Table 1.** Cluster Gene enrichment of combined bone marrow and spleen Treg dataset: differential expression of all genes per cluster against the rest of clusters, where measures of score, fold change and adjusted p-value are reported. Score measurement is calculated by SCANPY algorithm and uses a formula based on fold changes and p-values. The genes in each cluster are sorted based on their score.

**Supplementary Table 2.** CXCR4+ gene enrichment of combined bone marrow and spleen Treg dataset: list of all genes sorted by significance (SCANPY score) resulting from differential gene expression between CXCR4+ cells and the rest of cells in the dataset. Measurements of fold change and adjusted p-value are also reported. CXCR4+ cells are identified as described in the methods.

## Notes

**Disclosure of interest:** The authors report no conflict of interest.

### Competing Interest Statement

The authors have declared no competing interest.

